# Electron microscopy reveals toroidal shape of master neuronal cell differentiator REST – RE1-Silencing Transcription factor

**DOI:** 10.1101/2022.10.06.511111

**Authors:** Pavel Veverka, Tomáš Brom, Tomáš Janovič, Martin Stojaspal, Matyáš Pinkas, Jiří Nováček, Ctirad Hofr

## Abstract

The RE1-Silencing Transcription factor (REST) is essential for neuronal differentiation. Here, we report the first 18.5-angstrom electron microscopy structure of human REST. The refined electron map suggests that REST forms a torus that can accommodate DNA double-helix in the central hole. Additionally, we quantitatively described REST binding to the canonical DNA sequence of the neuron-restrictive silencer element. We developed protocols for the expression and purification of full-length REST and the shortened variant REST-N62 produced by alternative splicing. We tested the mutual interaction of full-length REST and the splicing variant REST-N62. Revealed structure-function relationships of master neuronal repressor REST will allow finding new biological ways of prevention and treatment of neurodegenerative disorders and diseases.

## Introduction

Human transcription factors control gene expression in each stage of cell development (1). Neuronal cell development is tightly regulated by the RE1-silencing transcription factor that binds a 21 bp silencer element (RE1) in the type II sodium channel gene (2). REST silences the expression of type II reporter genes in neuronal cells (3). The RE1-silencing transcription factor is called REST, but it is also known as NRSF – neuron-restrictive silencer factor (4). Activated REST ensures normal aging in human cortical and hippocampal neurons, probably due to the prevention of oxidative stress and amyloid β-protein toxicity (5).

REST is widely expressed during embryogenesis and plays a critical role in the final stage of neuronal differentiation (3,4). Additionally, REST is expressed in differentiated neurons during the critical time of enhanced sensitivity to plasticity in brain development (6). REST fine-tunes genes controlling synaptic plasticity (7). Furthermore, REST protects neurons by suppressing genes involved in the death of aging neurons (5).

REST expression changes are associated with neurodegenerative disorders and diseases (6,8). REST derepression is involved in induced neuronal death, as ischemic insults derepress REST in neurons destined to die (9). REST is activated in selectively vulnerable mature hippocampal neurons in response to seizures (10). In Huntington’s disease, REST expression increases in the nuclei of striatal neurons (11). In aging neurons, loss of REST is associated with the onset of Alzheimer’s disease in humans (5).

The combination of bioinformatical and biochemical analyses identified 1892 putative coding and non-coding genes in the human genome that REST can target (12).

REST is a 1097 amino acid, 116 kDa protein that contains a total of nine zinc fingers (ZFs). A cluster of eight zinc fingers constitutes the zinc finger domain, which mediates specific recognition and binding to DNA (Figure 1a). A combination of EMSA and molecular dynamics experiments suggests that ZFs 3–8 are inserted into the major groove of DNA duplex. Each ZF recognizes a triplet of canonical base pairs (13).

**Figure 1.**
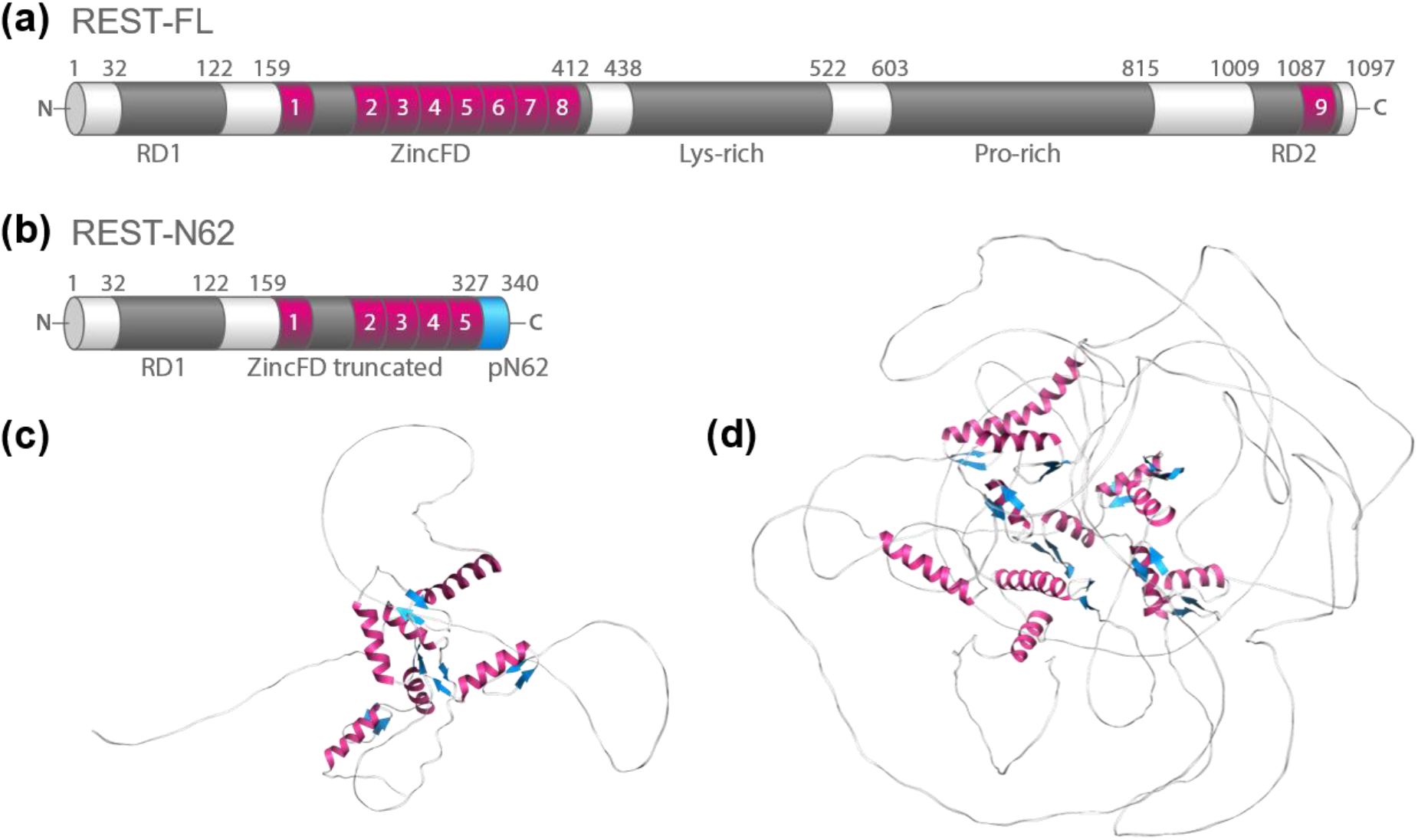
Expression variants of human REST – domain composition and structure prediction. (a) Full-length REST (REST-FL) comprises N-terminal repression domain 1 (RD1), Zinc finger domain (ZincFD) with eight zinc fingers mediating DNA binding, lysine-rich and proline-rich domains, C-terminal repression domain 2 (RD2) with the ninth zinc finger. Magenta regions depict zinc fingers. (b) REST-N62 is a splice variant of human REST that comprises only five zinc fingers and an extra thirteen amino-acid peptide insert. The cyan region represents the insert peptide N62 (pN62) that is generated by the alternative splicing of a neural-specific exon causing premature translation termination of truncated REST variant after inserting extra 62 nucleotides encoding 13 amino acids. Predicted structures of (c) REST-N62 ([AF-A0A087X1C2-F1](https://alphafold.ebi.ac.uk/entry/A0A087X1C2)) and (d) REST-FL ([AF-Q13127-F1](https://alphafold.ebi.ac.uk/entry/Q13127)) deposited in the AlphaFold Protein Structure Database. The helices of zinc fingers are shown in magenta; beta sheets are in cyan.

The function of REST as a transcriptional repressor is mediated by the N-terminal repressor domain (RD1) recruiting corepressor Sin3a/b that attracts histone deacetylases to remove acetyl groups from the core histones. The deacetylation of histones makes chromatin tightly packed, so the genes are inaccessible to the transcriptional machinery (14). Additionally, the C-terminal repressor domain (RD2) recruits corepressor complex CoREST, including histone deacetylases and methyltransferases. Methyltransferases induce the binding of Heterochromatin protein 1 to histones leading to the transcriptionally silent state of heterochromatin (15). Hence, REST causes gene repression through changes in the chromatin structure that prevent transcription (14,16).

Although human REST occurs in several isoforms (17), the contribution of various REST isoforms to neural induction is still under-investigated. For example, the most sequentially altered variant occurring specifically in human neurons is the REST-N62 isoform. REST-N62 (also known as REST4 or REST4-S3), which lacks four ZFs [6-9] and the C-terminal repressor domain, is generated by the alternative splicing of a neural-specific exon N located between exon IV and V (Figure 1b). The alternative splicing produces a shortened variant of REST that contains an extra 62 nucleotide insertion in the REST mRNA that encodes 13 amino acids and bears an in-frame stop codon that causes premature translation termination and synthesis of truncated REST variant (17).

Interestingly, REST-N62 has been shown to counteract the repressive role of full-length REST in the regulation of the target genes during neurogenesis (17–19). Recently, it has been suggested that REST-N62 binds full-length REST and the complex formation modulates REST function by changing the number of active REST molecules (20).

Here we addressed the following questions that are essential for further understanding the mechanism of REST function at the molecular level: 1. How to produce REST *in vitro* for functional and structural analyses? 2. What is the binding affinity of REST to the canonical DNA? 3. Does REST interact directly with the splicing variant REST-N62? 4. What is the structure of full-length REST? Despite being of undeniable importance, our understanding of REST function at the molecular level is still negligible. In this work, we successfully expressed and purified REST. We disclosed the first structure that helps us to understand REST spatial arrangement and provides implications for gene repression mechanisms in cells.

## Results

### Structure prediction revealed high ratio of intrinsically disordered regions

Initially, we analyzed the predicted structures of full-length REST (REST-FL) and the truncated variant produced by alternative splicing (REST-N62). We used the structures of REST-FL (Q13127), and REST-N62 (A0A087X1C2) deposited in the AlphaFold Protein Structure Database (21,22). To ensure the precise identification of the protein sequence and predicted models, we include unique UniProtKB and AlphaFold accession numbers, respectively.

The visualization of predictions of REST-FL ([AF-Q13127-F1](https://alphafold.ebi.ac.uk/entry/Q13127)), and REST-N62 ([AF-A0A087X1C2-F1](https://alphafold.ebi.ac.uk/entry/A0A087X1C2)) and shown in Figures 1 revealed central structured regions surrounded by intrinsically disordered regions (IDRs). The structured parts belong almost exclusively to the zinc finger domains. Each zinc finger domain consists of an alpha-helix and two associated antiparallel beta-sheets. The structural prediction suggests that the structured regions occupy 19% of the REST-FL sequence, while the content of structured regions in the variant REST-N62 is 30%, suggesting that REST-N62 might be more structurally stable than the full-length protein. Taken together, both REST-FL and the truncated isoform REST-N62 show high overall structural flexibility, which may contribute to the ability of REST to recruit various factors involved in chromatin structure arrangements.

### Establishing expression and purification procedures for recombinant REST-N62 and REST-FL

Expression and purification of the full-length REST are challenging mainly because of its large size. Additionally, the high frequency of IDRs and DNA binding capability make REST-FL an extremely challenging target for expression. Prediction studies indicated that the shorter variant REST-N62 might be more structurally stable than full-length REST. Despite our previous experience with preparing proteins, the purification procedure development for REST was challenging and lengthy. To prevent the scientific community from repeating unsuccessful ways of purification, we included a section in the discussion that describes the approaches we have tried out in our effort to purify REST-N62 and REST-FL.

After considering the reasonable length of REST-N62, we decided to express REST-N62 in *E. coli*, as bacteria usually produce relatively high amounts of recombinant proteins. We used an expression construct based on vector pl21 containing CL7 affinity tag, SUMO-tag with UlpI cleavage site, and the gene for REST-N62 (340 coding amino acids) followed by a stop codon. The protein was expressed in *E. coli* strain BL21-CodonPlus® (DE3)-RIPL in a TB medium.

To purify bacteria-expressed REST-N62, we first loaded processed supernatant of crude extract on the ultra-high affinity chromatography column Im7, which strongly bound CL7-tagged REST-N62 (Figure 2a, and Supplementary Figure S4a). In the second stage, after we removed the tags from REST-N62 by cleavage with UlpI protease, we loaded the protein and protease mixture on the His-trap column to separate mainly protease and chaperones from the REST-N62 (Figures 2a and 2b). In the final polishing step, we achieved high purity of REST-N62 using size exclusion chromatography that separated the remaining UlpI protease (Figures 2c and 2d).

**Figure 2:**
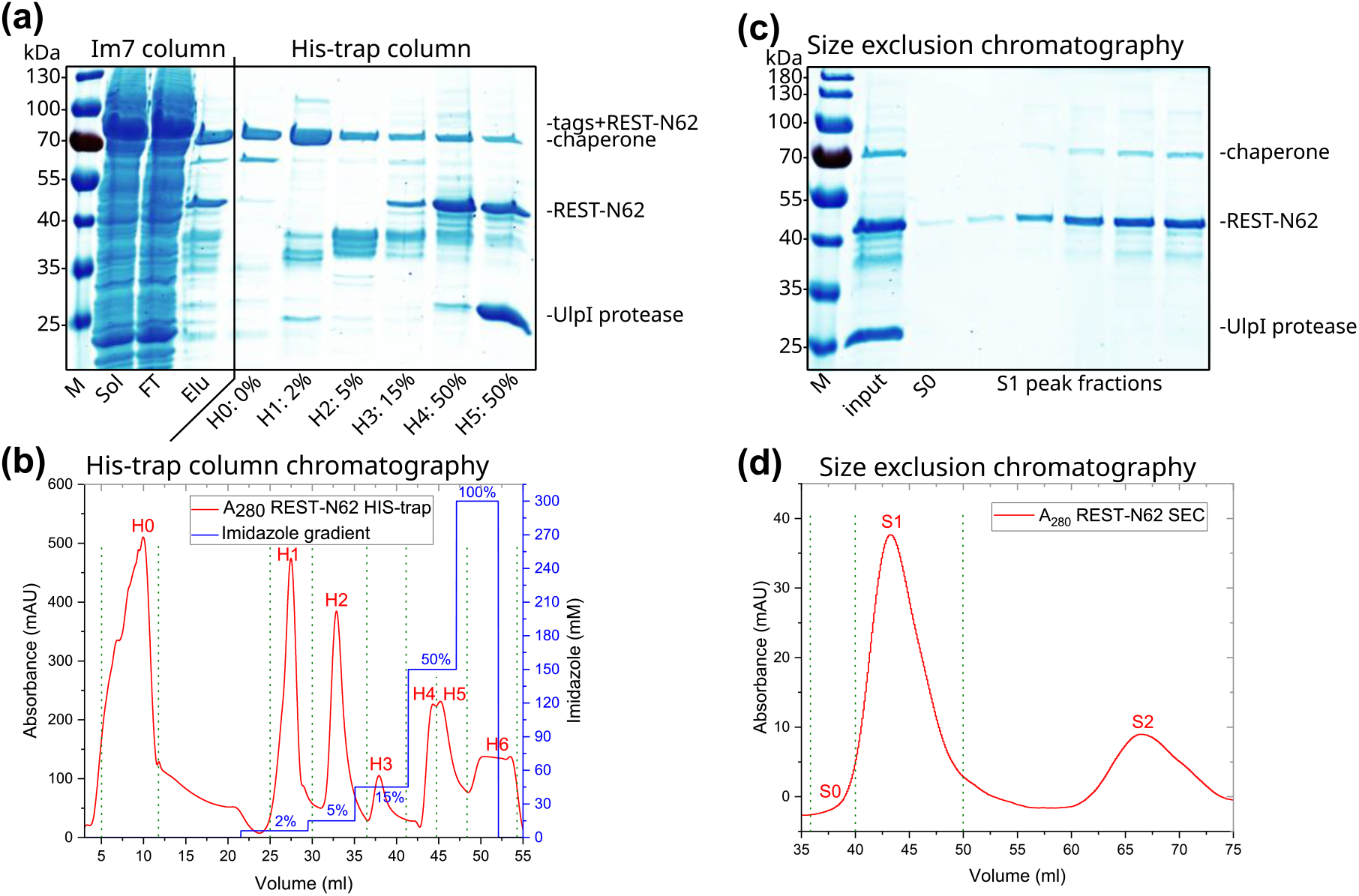
REST-N62 purification. (a) Analysis of Im7 and His-trap purification steps using 12% SDS-PAGE. M – protein molecular weight marker; Sol – soluble fraction of crude extract; FT – flow-through fraction; Elu – collected elution fraction from Im7 column after cleavage. His-trap column elution samples are denoted with peak number and percent of imidazole gradient: H0 – a fraction with washed-out chaperone without imidazole. H1, H2, and H3 – fractions with 2%, 5%, and 15% of imidazole, respectively. H4 and H5 – fractions with REST-N62 and UlpI protease, pooled and used as input for size exclusion chromatography. (b) His-trap column chromatography traces. His-trap column separated the chaperone and low molecular weight contaminations from REST-N62. Fraction marks correspond to the fractions analyzed in (a). Fractions H4 and H5 contained REST-N62, and UlpI protease were pooled. Imidazole absorbance is projected to the absorbance trace according to the imidazole concentration (H6 fraction). Green marks denominate collection ranges. (c) Analysis of size exclusion chromatography fractions by 12% SDS-PAGE. M – protein molecular weight marker. Input – pooled H4 and H5 fractions from previous His-trap purification. S0 – pre-peak fraction, S1 – collected peak fractions. (d) Size exclusion chromatogram of REST-N62. SEC separated protease from REST-N62. S0 – pre-peak fraction, S1 – main peak fractions with REST-N62, S2 – peak corresponding to UlpI protease. Green marks denominate collection ranges.

After extensive trials, we decided to express REST-FL in insect cells because insect cells can easily express proteins of size over 100 kDa. An additional advantage of the expression in the eukaryotic system is the maintenance of posttranslational modifications, including phosphorylation. REST-FL can be phosphorylated at phospho-serines in a conserved phosphodegron – a short linear motif that can interact with ubiquitin ligases upon phosphorylation. The phosphodegron is located in the C-terminus of REST-FL (23). We also tested the expression of REST-FL using the mammalian HEK293T cell line system. However, when we detected unwanted DNA contamination in REST-FL extracted from HEK cells (Supplementary Figure S2), we decided to express REST-FL in insect cells

We have designed over thirty vector constructs for insect cell expression of REST-FL. Eventually, we found the vector construct that was the most efficient for REST-FL expression. The selected pFASTBAC1 expression vector construct contains inserts in the following order: MBP-tag, 6xHis, S-tag, CL7 affinity tag, SrtA cleavage site, REST-FL gene (1097 coding amino acids). The expression vector construct was introduced into Sf9 insect cells. The Bac-to-Bac insect cell expression system was used to generate a baculovirus that expressed REST-FL.

To purify insect-cell expressed REST-FL, in the first step, we loaded processed supernatant of crude extract on the ultra-high affinity chromatography column Im7 that specifically bound CL7-tagged REST-FL (Figure 3a and Supplementary Figure S4b). In the second step, after cleavage with SrtA protease, we loaded the mixture of REST-FL and protease on a His-trap column to separate the protease and purified REST-FL (Figures 3a and 3b).

**Figure 3:**
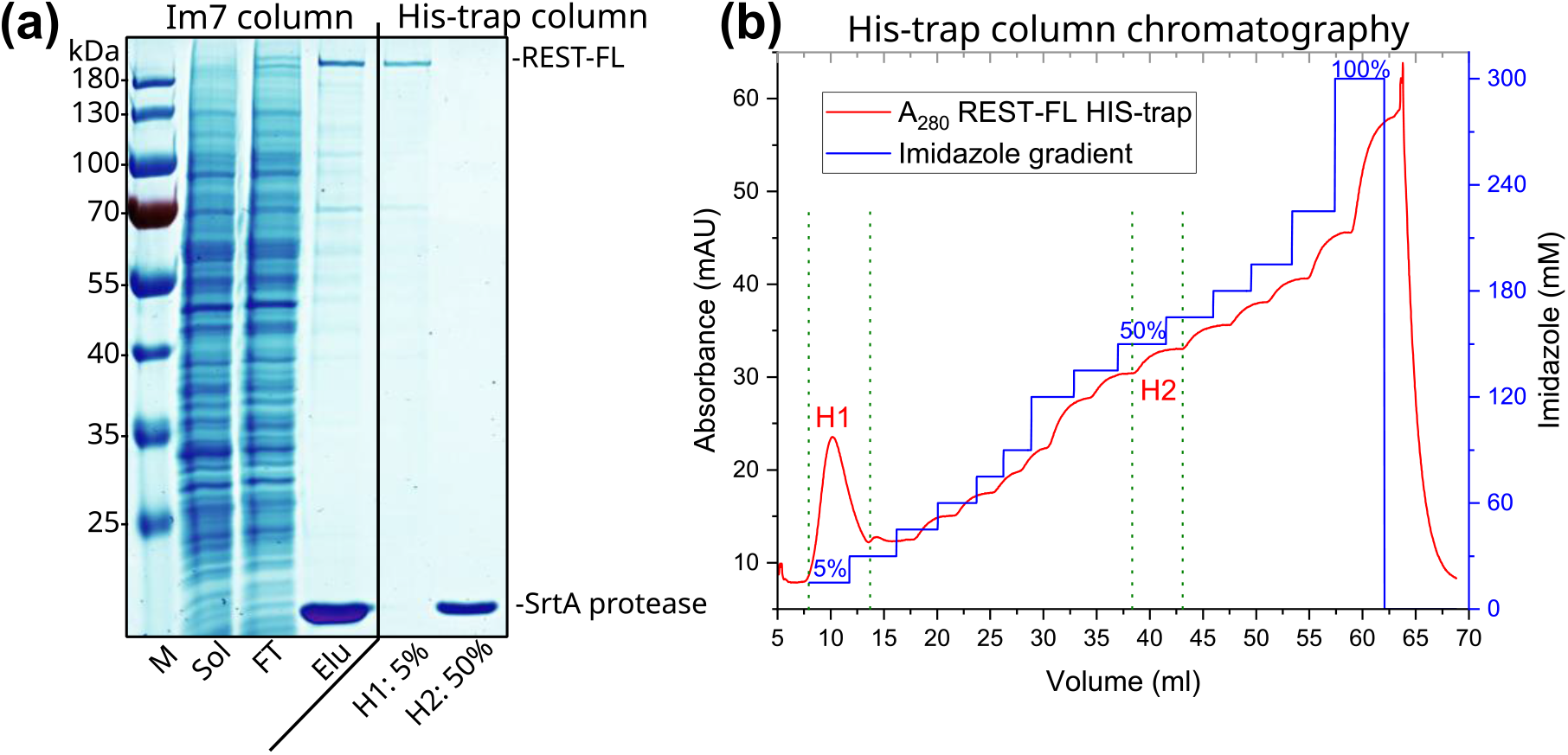
REST-FL purification. (a) Analysis of Im7 and His-trap purification steps using 10% SDS-PAGE. M – protein molecular weight marker; Sol – soluble fraction of crude extract; FT – flow-through fraction; Elu – collected elution fraction from Im7 column after cleavage. His-trap column elution samples are denoted with peak number and percent of imidazole gradient: H1 (5% imidazole) contains REST-FL, and H5 (50% imidazole) contains SrtA protease. (b) His-trap column chromatography traces. His-trap column separated the protease and low molecular weight contaminations from REST-FL. Fraction marks correspond to the fractions analyzed in (a). Fraction H1 contained REST-FL. H2 fraction corresponds to SrtA protease. Imidazole absorbance is projected to the absorbance trace according to the imidazole concentration. Green marks denominate collection ranges.

We have described the expression and purification of REST-N62 and REST-FL in detail in the Materials and methods section.

### Quality, homogeneity, and DNA content of recombinant REST

The quality of REST-FL and REST-N62 was analyzed by SDS-PAGE (Figure 4). The purity values REST-N62: 95%, and REST-FL: 85% determined by densitometry analysis were sufficient for subsequent experiments. We confirmed the identity of REST-N62 by MALDI-TOF mass spectrometry. Experimentally measured molecular weight (38 370 Da) was comparable to calculated molecular weight (38 379 Da), suggesting that REST-N62 was prepared without extra tags, only with remaining glycine and serine residues at the N-terminus after UlpI protease cleavage (Supplementary Figure S1).

**Figure 4:**
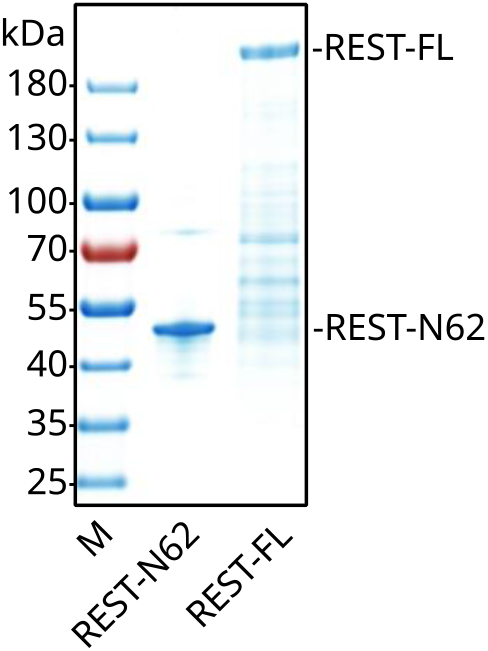
Final purity of REST constructs. Analysis of REST-FL and REST-N62 after purification using gradient SDS-PAGE. M – protein molecular weight marker.

In the case of REST-FL, MALDI-TOF mass spectrometry provided suppressed signal because of the presence of molecules with lower molecular weight than REST-FL. Therefore, we used MALDI MS/MS analysis of the tryptically digested band extracted from SDS-PAGE to identify REST-FL. The MS/MS spectrum was compared to the sequence from UniProt database accession number Q13127. The MASCOT score of 847 and 23% sequence coverage of MS/MS analysis confirmed that the electrophoretic band with the expected molecular weight is exclusively formed by REST-FL.

As REST-FL serves as a transcription factor with high affinity to DNA, we measured the possible content of genomic DNA in the purified proteins before performing quantitative DNA studies.

We compared the DNA content in the REST-FL samples expressed in mammalian and insect cells by combining UV spectroscopy and gel electrophoresis. UV spectroscopy revealed an absorbance ratio A260/A280 1.62 ± 0.08 for REST-FL expressed in mammalian cells and 0.49 ± 0.07 for REST-FL expressed in insect cells. The A260/A280 ratio higher than 0.6 suggested that genomic DNA contaminated REST-FL isolated from mammalian cells. On the other hand, REST-FL isolated from insect cells showed the ratio corresponding to the absence of DNA. To confirm that REST-FL isolated from insect cells is DNA-free, we separated REST-FL and REST-N62 on an agarose gel (Supplementary Figure S2). No DNA traces were found in REST-FL from insect cells.

DNA-free REST-FL from insect cells was used for subsequent binding and structural studies. REST-N62 could be produced in bacteria in significantly higher amounts than REST-FL in insect cells. Hence, we used more sample demanding methods to evaluate REST-N62 homogeneity and secondary structure. Dynamic light scattering (DLS) revealed that REST-N62 is homogenous with an expected average radius of 10 nm without significant aggregations (Supplementary Figure S3a). Minor content of a chaperone in the sample (Figure 4) may contribute to the observed peak at around a 60 nm radius. Moreover, the proper folding and secondary structure composition of REST-N62 were assessed by far-UV circular dichroism (CD). The recorded CD spectrum suggested that REST-N62 is well-folded with prevalently alpha-helical arrangements of structured regions (Supplementary Figure S3b). The observed prevalence of alpha-helices in the secondary structure is in accordance with the prediction of the REST-N62 structure (Figure 1c). In summary, our analyses confirmed that recombinantly prepared REST-FL and REST-N62 meet the requirements of purity, homogeneity, and DNA absence for further studies.

### Purified REST-FL binds NRSE with high affinity

We quantified the affinity of REST-FL and REST-N62 to canonical DNA using fluorescence anisotropy. Both REST variants were allowed to bind to an oligonucleotide duplex containing the NRSE canonical sequence (21bp) labeled by AlexaFluor 488 (Figure 5). We found that purified REST-FL binds NRSE with a high-nanomolar affinity. The analysis of the data using a one-site binding model revealed dissociation constant K_D_ = 5.7 ± 0.6 nM. Conversely, REST-N62 showed no binding capability (Figure 5) in the same concentration range. We show that purified REST-FL actively binds canonical DNA with a nanomolar dissociation constant.

**Figure 5.**
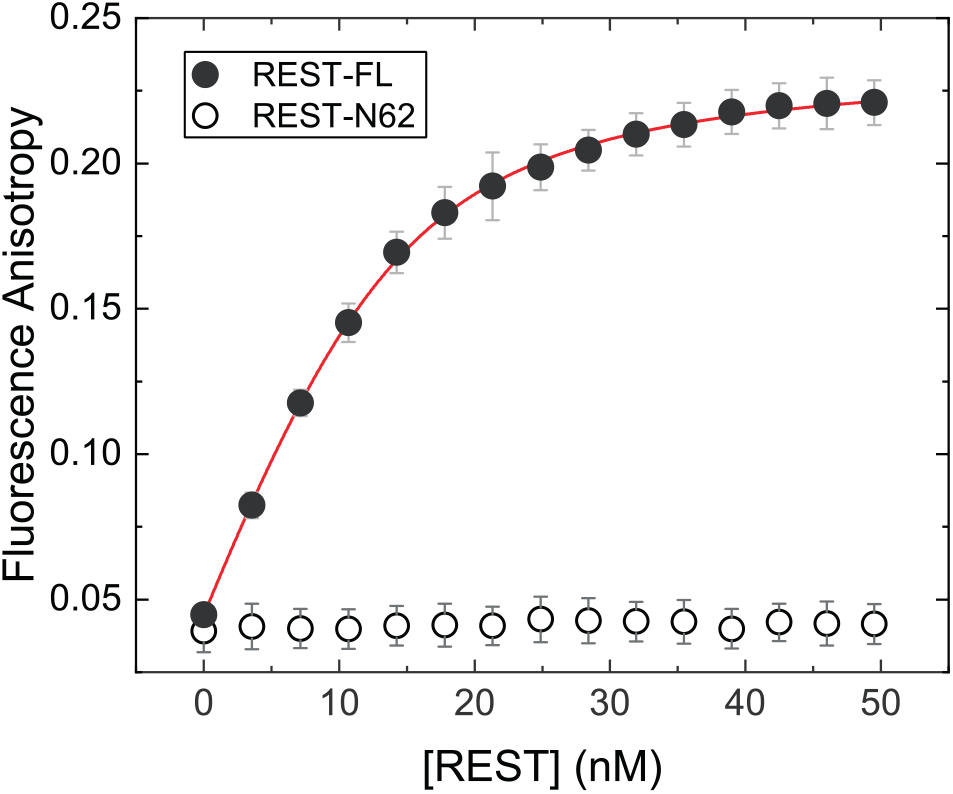
REST-FL and REST-N62 binding to NRSE DNA recognition sequence. REST variants were allowed to bind oligonucleotide duplex containing NRSE canonical sequence labeled by AlexaFluor 488 (7.5 nM). The dissociation constant KD was determined by non-linear fitting using a one-site binding model (red). REST-N62 showed no significant DNA binding affinity; no fitting model was applied. Error bars represent s.d., n = 3 independent measurements.

### Coimmunoprecipitation detected no interaction between REST-N62 and REST-FL

We wondered if REST-N62 and REST-FL would interact together. The mutual binding of REST-N62 and REST-FL was hypothesized in previous studies (18) and later suggested as a regulatory mechanism (20). To test the hypothesis of mutual binding of REST-N62 and REST-FL, we performed coimmunoprecipitation (CoIP) of overexpressed proteins in HEK293T cells, followed by western blotting (WB) analysis. We immunoprecipitated FLAG-tagged REST-N62 using M2 FLAG magnetic beads to detect binding to REST-FL (Figure 6). Coimmunoprecipitation results showed no detectable mutual interaction of REST-N62 and REST-FL under our conditions. Hence, our data suggest a low possibility that the shorter isoform REST-N62 can interfere with and modulate the silencing activity of REST-FL.

**Figure 6:**
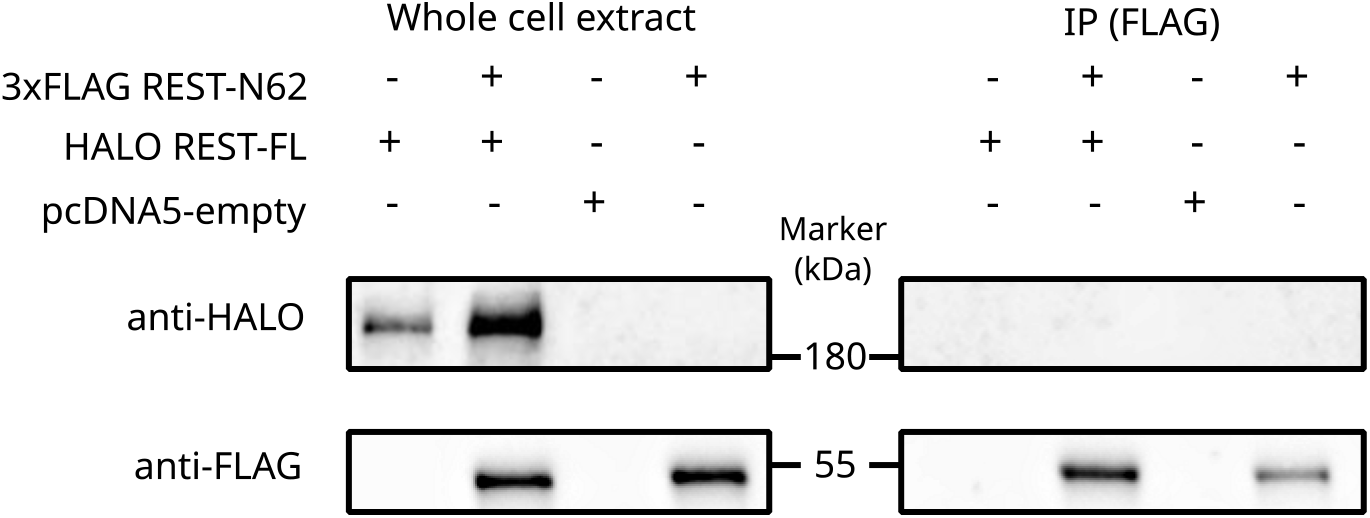
Coimmunoprecipitation of REST-FL and REST-N62 showed no mutual interaction. FLAG-tagged REST-N62, and HALO-tagged REST-FL variants were transiently co-expressed in HEK293T cells. REST variants were precipitated using M2-FLAG magnetic beads. Precipitates were analyzed by western blotting with corresponding antibodies.

### Negative stain electron microscopy reveals the toroidal structure of REST-FL

We solved the structure of purified recombinant human REST-FL at 18.5 Å resolution using negative stain electron microscopy. The subsequent 3D reconstruction revealed a toroidal structure of REST-FL. The reconstructed doughnut-like shape has an outer diameter of 14 nm and an inner diameter of 5 nm. The side view shows that the height of the toroid is 9 nm (Figure 7c).

**Figure 7:**
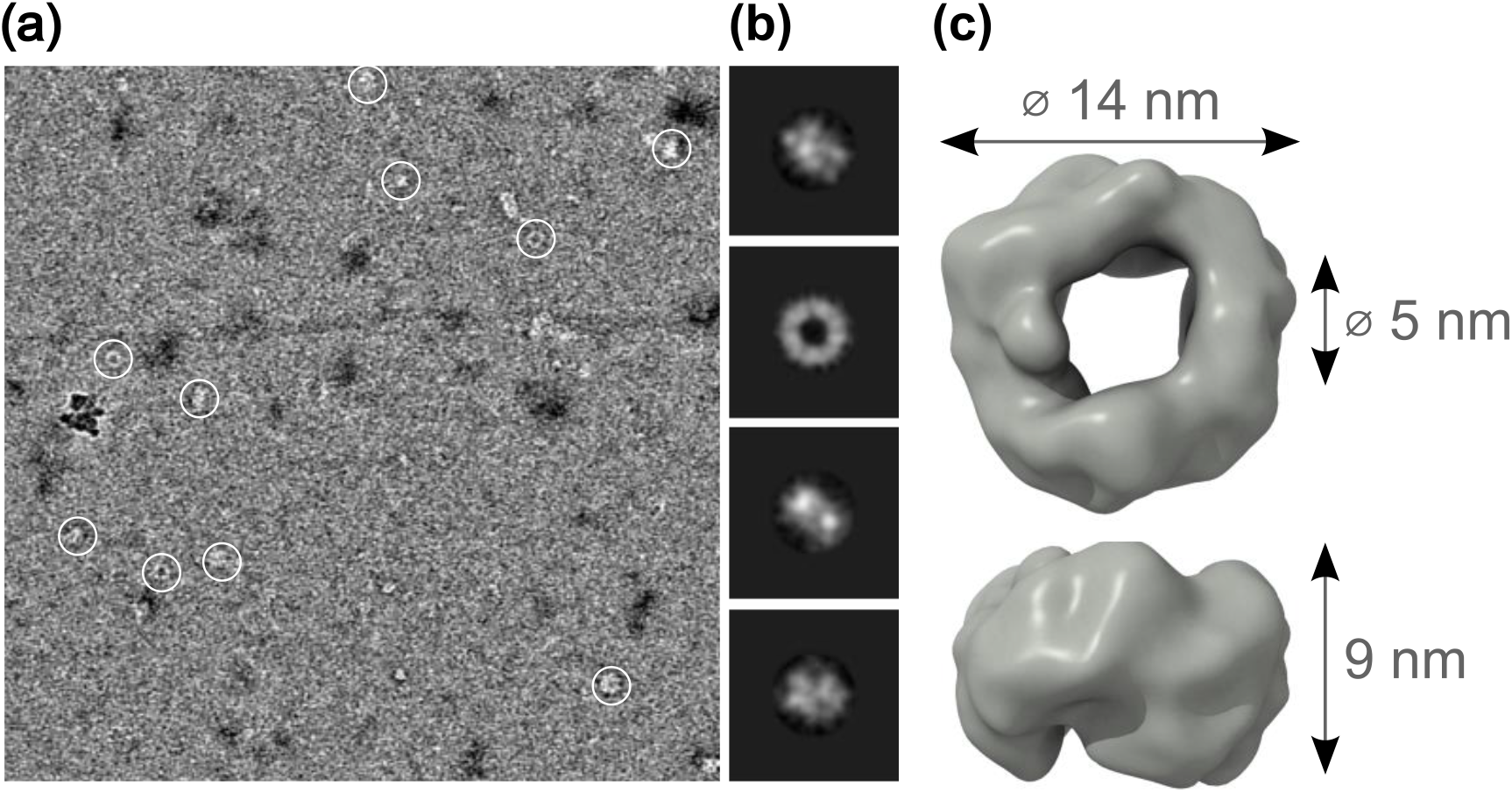
Negative stain electron microscopy of REST-FL. (a) Representative electron micrograph of REST-FL stained with 2% uranyl acetate. Protein particles are highlighted by white circles. (b) Representative results from the reference-free 2D classification of the REST particles. (c) 3D model of REST at 18.5 Å resolution (EMD-15858). Two different views of the 3D reconstruction rotated by 90° around a horizontal axis. The arrows indicate the size of the REST structure. The outer diameter of the model is approximately 14 nm, the diameter of the inner pore is approximately 5 nm, and the height is approximately 9 nm.

From the 2D class of top view, it might appear that the toroidal shape of REST-FL may adopt a multimer symmetry. Based on the prediction of the REST-FL monomer (Figure 1d), we selected the partially structured N-terminal part (150-425 aa) comprising eight zinc fingers and predicted the dimer, trimer, tetramer, pentamer, and hexamer using AlphaFold v2.1.0 multimer tool (22). The predicted models showed no mutual interaction of monomeric subunits nor adopted any defined multiunit arrangement of REST-FL through the structured regions.

After successful analysis by negative staining, we carried out extensive cryo-EM experiments. REST-FL grids were prepared with storage buffer containing 2.5 % glycerol. Unfortunately, the glycerol content strongly affected the ice thickness, ice formation and negatively impacted the particle contrast. We attempted to prepare cryo-grids with a broad range of lower glycerol concentrations. In all cases, we observed substantially more pronounced aggregation of protein mass on the cryo-grids compared to negative staining. Thus, we preferentially concentrated our effort on precisely analyzing negative stain electron microscopy grids.

## Discussion

We revealed the electron microscopy structure of master neuronal cell differentiator RE1-Silencing Transcription factor (REST). Additionally, we developed protocols for the expression and purification of full-length REST and its splicing variant REST-N62. We quantified REST binding to canonical NRSE sequence. Furthermore, our findings suggest no mutual direct binding of REST-FL and REST-N62. Our studies opened new horizons to extensive studies with full-length REST and its splicing variant REST-N62. We thoroughly discuss our findings and their implications below. We divided the discussion into three sections covering the preparation, binding, and structure of REST.

### Challenges in expression and purifications – dead ends to avoid

This section is dedicated to all colleagues scientists trying to optimize REST production. We described our experimental approaches, including unsuccessful ones, to avoid repeating the ineffective ways of REST-FL and REST-N62 production.

When we were preparing expression vector constructs containing the REST-FL gene, we observed that the gene comprises several palindromic sequences. The repetitive sequences, along with frequent off-frame hexahistidine sequences, might cause cloning difficulties.

We tested many possible combinations of tags for expression, solubility improvement, and affinity purification, including 6xHis, S-tag, GFP, HALO, SUMOstar, 3xFLAG, thioredoxin, CL7, and MBP. Surprisingly, we found that HRV 3C protease, which we use preferentially to cleave expression tags, cleaves REST-FL nonspecifically at amino acid region 654-661: MEVVQEGP. Therefore, the initial use of HRV 3C in our expression trials caused a low yield of full-length REST.

We consequently tried other proteases such as TEV, WELQut, UlpI-star, and SrtA (SortaseA), with the last being the best for our purposes. Additionally, we tried different purification sorbents compatible with the previously mentioned affinity purification tags and corresponding purification procedures.

Initially, we tried REST-FL expression in mammalian cells HEK293T, but we observed that the purified protein contained genomic DNA (Supplementary Figure S2). When we added benzonase at the initial stage of purification, we observed the low DNA binding ability of REST-FL to NRSE DNA sequence (data not shown). Our inability to produce active REST-FL in mammalian cells drove our attention to the insect cell expression system.

During the optimization of REST-FL production in insect cells, we tested several bac-to-bac and multibac vectors with different promotors such as polyhedrin, p10, hr5, and IE1 with corresponding terminators, and different competent cells for bacmid production such as DH10Bac, DH10MultiBac, and DH10EmBacY, before we found the optimal combination listed in the Materials and methods.

The choice of the final purification step is essential for the resulting REST-FL activity. As REST-FL contains nine zinc-finger domains that require Zinc cations for structure stabilization, we tested if charging the immobilized metal affinity column with Ni^2+^ or Zn^2+^ ions affects REST-FL activity. We observed no significant effect of the ion substitution on the DNA binding activity or structure of REST-FL (data not shown).

In general, we found that the purification of recombinant REST-FL produced in insect cells is suitable for *in vitro* analyses, including electron microscopy, because insect cells contain no NRSE DNA sequence that could naturally bind and block the activity of REST-FL.

REST-N62 (38.2 kDa) is significantly smaller than REST-FL (121.9 kDa), so we found REST-N62 expression and purification less complicated. We optimized the preparation of REST-N62 in *E. coli* via different expression setups, including various solubility and purification tags – 6xHis, S-tag, 3xFLAG, thioredoxin, Sumo, and CL7 together with proteases HRV 3C or UlpI. Thioredoxin usually improves solubility; however, in this case, the thioredoxin tag induced immediate precipitation of REST-N62 after the lysate sonication.

The optimal combination of temperature and aeration during REST-N62 expression was essential for proper protein production without excessive chaperone expression.

For example, when we used a different media:air volume ratio than the optimal 0.5L TB medium in 2L Erlenmeyer’s flasks, the DnaK chaperone from *E. coli* was overexpressed. Similarly, when we reduced the temperature from 37 °C to 20 °C before the induction of expression, we discovered that cooling must be finished within one hour to ensure that the minimum amount of chaperone DnaK is produced. DnaK presence in samples was confirmed by MS analysis (data not shown). This chaperone can be observed on SDS-PAGE gel (Figure 2) at the level of 70 kDa protein marker.

Despite all before mentioned challenges, we successfully optimized protocols to recombinantly produce active REST-FL and well-folded REST-N62 *in vitro*. The final and optimal expression and purification protocols are in the Materials and methods section.

### REST-FL binds avidly to DNA but is reluctant to interact with REST-N62

As REST is a transcription factor that binds specifically canonical sites on DNA, we might expect that REST-FL shows DNA binding affinity similar to proteins that bind and recognize specific DNA sequences (24,25). Human transcription factors that can interfere with genomic DNA usually bind with K_D_ in the nanomolar range. The observed REST-FL K_D_ value is in the range that is common for human transcription factors (26).

On the contrary, REST-N62 showed no DNA binding capability (Figure 5) in the same concentration range as REST-FL. Even though REST-N62 shares the same 1-5 zinc finger motifs on its N-terminus (Figure 1), the missing zinc fingers 6-9 seems to be the critical part for high-affinity binding to DNA. Indeed, Tang et al. showed that zinc fingers 7, and 8 contribute significantly to recognizing DNA with NRSE sequence. According to previously published results of combined electromobility shift assay and molecular dynamics with zinc fingers 1-5, mainly zinc fingers 4 and 5 contribute to DNA binding (13). Hence, REST-N62 may still bind DNA through zinc fingers 4 and 5 if the spatial orientation is preserved as in REST-FL. The binding via only two zinc fingers may explain why we observed no significant fluorescence anisotropy increase during the addition of REST-N62 to the canonical NRSE DNA (Figure 5). Additionally, we tested the binding of REST-N62 to NRSE DNA motif at concentrations over 1.2 mM (data not shown), but we observed no significant binding of REST-N62 either. However, we cannot exclude that REST-N62 may still bind NRSE DNA with substantially lower affinity with a corresponding K_D_ value several orders of magnitude higher than full-length REST.

Finally, to investigate the possible mechanism of REST regulation on a molecular level, we tested the previously suggested hypothesis that REST-N62 isoform binds REST-FL to block the ability of REST to interact with DNA (20). In our conditions, coimmunoprecipitation experiments in mammalian cell lysate showed no detectable interaction between human REST-N62 and REST-FL (Figure 6). We suggest that the splicing variant REST-N62 does not interact directly with full-length REST.

### The toroidal structure of REST allows central DNA binding that is common for transcription factors

Here we described the first structure of full-length human REST using negative staining electron microscopy.

The 3D reconstruction suggests a toroidal shape. The doughnut-like shape is typical among proteins that serve as molecular machines participating in DNA replication and RNA transcription (27). For example, β-clamp polymerase processivity factor (PDB: 3BEP) and Proliferating cell nuclear antigen (PCNA) (PDB: 1AXC) also possess a doughnut-like structural arrangement. The inner diameter of REST central pore (5 nm) is comparable to the pore sizes of other toroidal DNA binding proteins (~4 nm) (Figure 8). The initial structure prediction of REST-FL by AlphaFold revealed the prevailing content of flexible and intrinsically disordered regions. The flexible IDRs of REST-FL might become rigid upon binding to DNA by zinc fingers or upon binding to other accessory factors.

**Figure 8:**
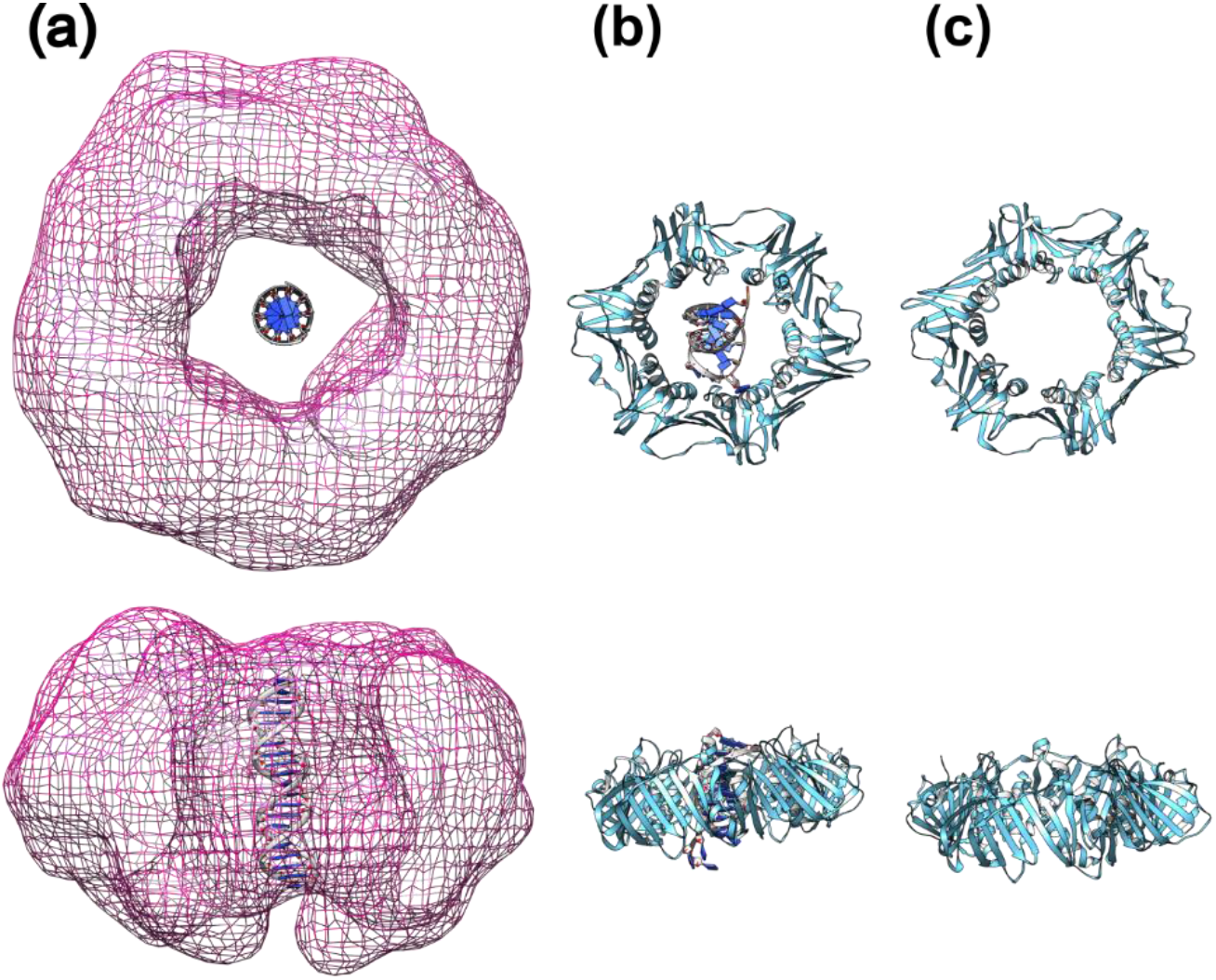
Putative model of NRSE embedded inside REST-FL envelope compared with toroidal β-clamp polymerase processivity factor. (a) Mesh model of REST-FL obtained by negatively stained electron microscopy with the projected NRSE sequence (EMD-15858). (b) β-clamp polymerase processivity factor cartoon model with bound DNA in the inner pore (PDB: 3BEP) and (c) without DNA. Lower images represent the 90° rotation of models alongside the x-axis.

Recently, Li et al. provided several modes of functional motions that may modulate the structure of toroidal systems upon binding of substrates. The toroidal motion modes include cooperative saddle-like bending, rolling, counter rolling, stretching and contraction, breathing, and clamshell-like motions (27). The predicted high dynamics of toroidal protein complexes enable toroids to undergo several motions simultaneously, resulting in harboring specific nucleic acid.

Our agarose gel analysis allows us to consider that REST-FL produced in insect cells contains no or a minimal amount of bound DNA (Supplementary Figure S2). Then electron microscopic reconstruction might represent the relaxed form of full-length REST structure without DNA and with a wide open central pore. We hypothesize that the pore size might be reduced upon REST binding to DNA due to the dynamic modes of motion that allow tightening of the pore and DNA stabilization within the toroid.

The toroidal complexes often bind DNA as oligomers. For instance, PCNA forms a trimer; the β-clamp polymerase processivity factor acts as a dimer. Thus, we may speculate that REST is capable of forming organized multimers. We have no data available to validate the monomeric or the multimeric arrangement of REST. We employed AlphaFold v2.1.0 multimer tool (22) to predict dimeric, trimeric, tetra-, penta- and hexameric structures of REST-FL or the most structured part of REST comprising eight zinc finger domains with no success.

Regardless of the monomeric or multimeric arrangement, we speculate that the zinc finger domain may be arranged facing the inner central pore of the toroidal structure allowing zinc fingers to access and “read” the DNA sequence. Figure 8 shows our putative model of the EM structure of REST with the canonical NRSE DNA projected in the central hole (Figure 8a). Remarkably, the EM structure of REST-FL easily accommodates the whole 21 base pairs of NRSE DNA sequence, as the 90° rotated model demonstrates in the lower part of Figure 8a.

To compare REST with other transcription factors, we included the known structure of toroidal β-clamp polymerase processivity factor with and without DNA (Figure 8b and 8c). We speculate that the unstructured interacting regions RD1 and RD2 might be on the outer edge, ready to interact with REST cofactors and other binding partners. Notably, the revealed toroidal arrangement of REST allows simultaneous binding of canonical DNA and recruitment of multiple proteins.

Transactivation domains of transcription factors are often intrinsically disordered and acquire transient structure only upon binding of a partner. Additionally, the IDRs may enable REST to interact with more than two hundred different partners (28). Specific IDRs are Lysine-rich and Proline-rich regions that could function as scaffolds for the toroidal structure. Moreover, the structure formation could be affected by posttranslational modifications.

The phosphorylation of the phosphodegron region causes the increase of local negative charge that may attract positively charged parts of REST or other interacting partners. Thus, the phosphodegron might contribute to the stable tertiary structure of REST and the long-distance recruitment of partners. The specific posttranslational modifications of IDRs might be why AlphaFold failed to predict the structure of REST.

The structure of REST determines how effective finding and binding the NRSE site on DNA is in the initial step of REST-mediated gene repression. The gene silencing process is a complex combination of successive steps, including interactions with cofactors and other binding factors.

Two binding partners of REST - Sin3a/b (29) and CoREST (30) play critical roles in the recruitment of chromatin modifying enzymes HDAC1/2, BRG1, LSDS1, and G9, which affect chromatin condensation and thus inhibit gene expression.

The predicted structures in the AlphaFold database show that REST-interacting regions of Sin3a (205-480 aa) and Sin3b (59-105 aa) are mostly unstructured. Similarly, CoREST is mainly flexible, and the only structured region is dedicated to recruiting LSD1 histone demethylase (31). Generally, the interaction between two IDRs is preferred over the interaction of two structured regions (32). Nevertheless, REST could interact with its binding partners via IDRs that are arranged to organized structures only upon complexation. The formation of a hydrophobic helix has been described for binding the N-terminal RD1 domain of REST with PAH1 domain of Sin3b (33). Consequently, even Sin3a and CoREST interactions could be mediated by IDRs that adopt well-defined structures upon binding.

We might consider several ways how REST could be regulated on a cellular level. Firstly, REST complexation with partners may initiate allosteric changes of IDRs that may enhance REST affinity to the corresponding DNA sequence or release REST from DNA. Secondly, the REST function could be modulated by transporting REST from the nucleus to the cytoplasm. The example of targeted REST transport provides Htt protein which sequesters REST in the cytoplasm and thereby enables neuronal survival, neuronal plasticity, and dendritic growth (34). Additionally, the most investigated way of hw can be REST level regulated is the path length including phosphorylation, ubiquitination, and subsequent degradation by the proteasome. Probably, REST is phosphorylated by Casein Kinase 1, which makes REST available for subsequent ubiquitination. E3 ubiquitin-protein ligase β-TrCP adds ubiquitin units at the phosphodegron site of REST. Hence, structural changes induced by phosphorylation make the ubiquitination site on REST accessible for enzymes (35). Finally, proteins that sterically block REST against ubiquitination might modulate the REST function. For instance, telomeric protein TRF2 has been found to prevent REST ubiquitination (36).

Our results serve as a solid base for further investigations of REST to answer the questions that appeared during our studies. Future research should address the REST structure in complex with canonical and noncanonical NRSE DNA. Additionally, the stoichiometry of REST arrangement with and without DNA should be determined. Finally, subsequent studies should describe REST interactions with cofactors essential for regulating gene repression.

## Materials and methods

### The cloning, expression, and purification of REST-N62

REST-N62, also known as REST4-S3 is a truncated variant of REST-FL achieved via alternative splicing – after Serine 327 with additional 13 amino acids: VGYGYHLVIFTRV followed by a stop codon (sequence with accession number A0A087X1C2).

The source of the REST-N62 sequence was a plasmid obtained from Addgene (#41903). Recombinant expression cassette contained inserts in the following order: CL7 Affinity Tag, SUMO-tag with UlpI cleavage site, REST-N62 (340 coding amino acids), stop codon. We found out that REST-N62 purification is baffled with the thioredoxine tag in the original pl21 plasmid, so we decided to delete thioredoxine and the short linker between CL7 and SUMO tags. Plasmid pl21 (TriAltus) was truncated by digestion with restriction enzymes XbaI and XhoI (New England Biolabs). To complete the recombinant assembly reaction NEBuilder® HiFi DNA Assembly kit (New England Biolabs) was used with the combination of three different inserts produced by PCR: the first insert contained the original pl21 vector part – the non-coding sequence before the start codon, followed by the coding sequence of CL Affinity Tag; the second insert encoded SUMO-tag; the third insert contained the REST-N62 coding sequence with flanking 13 extra amino acids followed by a stop codon. All PCR products and digested plasmid were separated by agarose gel electrophoresis and extracted from the gel with QIAquick® Gel Extraction Kit (Qiagen). After assembly, the expression vector construct was transformed into *E. coli* One Shot™ TOP10 (Invitrogen). The final construct was confirmed by sequencing, and the plasmid with the coding REST-N62 sequence was transformed into *E. coli* expression strain BL21-CodonPlus® (DE3)-RIPL (Agilent).

Expression was carried out in 0.5L of Terrific Broth (TB) medium containing 50 μg·mL^-1^ kanamycin and 34 μg·mL^-1^ chloramphenicol in 2L Erlenmeyer’s flask to ensure good aeration. The preheated medium was inoculated with 10 mL of overnight grown BL21-(DE3)-RIPL cells. Flasks were shaken in orbital shaker New Brunswick™ Innova® 43/43R (Innova) at 160 rpm and 37 °C. After OD_600_ reached 1.0, the flasks were cooled down to 20 °C (typically one hour of cooling). Protein expression was induced by the addition of IPTG to the final concentration of 0.1 mM. The cell cultute was incubated overnight (at least 8 hours) and then collected by centrifugation (8 000 g, 8 min, 4 °C). Obtained pellet was immediately used for purification (without freezing). We usually performed expression in a total 3L TB medium and obtained around 30 g of cell pellet.

All the following purification steps were carried out with precooled buffers and instruments. Additionally, all following steps should be done at 4 °C. Fresh pellet kept on ice was resuspended in the lysis buffer (50 mM Sodium Phosphate, 500 mM NaCl, 5 % glycerol, 0.1 % Triton X-100, pH 8.0) with the addition of 0.8 mg·mL^-1^ of lysozyme, leupeptin 1.9 μg·mL^-1^ and pepstatin A 3.1 μg·mL^-1^ (80 mL of lysis buffer was used with typical 3L TB expression). The cell suspension was transferred into stainless steel container cooled on ice and sonicated for 12 minutes with a 30% amplitude, 1-second pulse-on, 1-second pulse-off with a 60-second process time cycle with a 15-second pause for extra cooling (Misonix Qsonica S-4000). The lysate was cleared by centrifugation at 20 000 g, 1 hour, 4 °C, and filtered by 0.45 μm Sterivex™ Filter Units (Merck Millipore).

REST-N62 was purified in three-step chromatography, carried out in a cold room at 4°C with a chromatography system (NGC™ Chromatography System Quest™ 10 Plus, Bio-Rad).

Ultra-high affinity chromatography column Im7 (TriAltus) was used to capture CL7-tagged protein. Typically, was used a 5 mL Im7 column and a loading flow rate of 0.5 mL·min^-1^. After loading, the column was washed with Im7 Wash buffer (50 mM Sodium Phosphate, 500 mM NaCl, 5 % glycerol, pH 8.0) until the absorbance at 280 nm was stabilized, followed by a second wash with SEC buffer (50 mM Sodium Phosphate, 150 mM NaCl, 2.5 % glycerol, pH 8.0). Cleavage was carried out by premixed 6xHis-UlpI SUMO protease (recombinantly prepared in our laboratory) to a final concentration of 35 μg·mL^-1^ in SEC buffer with the addition of DTT to the final concentration of 1 mM. The protease was loaded to the Im7 column in two steps divided by 20 min intermediate equilibration and followed up to a two-hour incubation. The cleaved protein was eluted at flow rate of 0.3 mL·min^-1^. After the first affinity purification step, we typically obtained 4 mg of the protein mixture, as shown in Figure 2a – Elu fraction.

Affinity chromatography with His-trap column (HisTrap™HP, Cytiva), typically 1 mL column volume, flow rate 1 mL·min^-1^, was employed in separating chaperone and other impurities from the sample. REST-N62 accommodates an unspecific binding to the column matrix. Elution was done in the 0-300 mM imidazole gradient in 50 mM Sodium Phosphate, 150 mM NaCl, 2.5 % glycerol, pH 8.0. Washing by 45 mM imidazole corresponding to 15% imidazole gradient removed impurities, as shown in SDS-PAGE Figure 2a. REST-N62 was eluted together with Ulp1 protease at 150 mM imidazole concentration (50% imidazole gradient) – typically total fraction volume 4 – 5 mL and 0.8 mg of total protein amount.

Size exclusion chromatography was employed to remove the remaining UlpI protease contaminant and change buffer composition. SEC column HiLoad® 16/600 Superdex® 75 prep grad (Cytiva) was equilibrated in two volumes of SEC buffer. The collected sample from His-trap selected fractions was centrifugated at 15 000 g, 10 minutes, 4 °C to ensure no protein aggregation before loading to the SEC column. SEC separation was carried out at a rate of 1 mL·min^-1^.

Collected samples from the first main peak were concentrated by ultrafiltration on 10K Amicon® Ultra-4 Centrifugal Filters Units (Merck Millipore) in a swinging-bucket rotor 4 000 g, 4 °C, until the concentration of protein was higher than 2 mg·mL^-1^. After concentrating, REST-N62 was frozen in liquid nitrogen and stored at −80 °C. Typical purification from 3L of TB media has a final yield of 0.3 mg of pure REST-N62. The final purity of REST-N62 is in Figure 4. Purification was carried out within one day to preserve protein activity.

### The cloning, expression, and purification of REST-FL

The source of the REST-FL sequence was a plasmid obtained from Addgene (#41903). Recombinant expression cassette contained inserts in the following order: start codon, MBP-tag, 6xHis, S-tag, CL7 Affinity Tag, short 90bp linker, SrtA site, REST-FL, stop codon. The pFASTBAC1 plasmid with polyhedrin promoter (Invitrogen) was linearized by digestion with restriction enzymes BamHI and HindIII (New England Biolabs).

To perform the recombinant assembly reaction, NEBuilder® HiFi DNA Assembly kit (New England Biolabs) was used with the combination of three different inserts produced by PCR: the first insert contained the MBP-tag, 6xHis, and S-tag from MBP-enhanced vector pTriEx4 (Novagen); the second insert contained CL Affinity Tag followed by a 90bp linker taken from pl21 plasmid (TriAltus); and the third contained the REST-FL coding sequence followed by a stop codon. All PCR products and digested plasmid were separated by agarose gel electrophoresis and extracted from the gel with QIAquick® Gel Extraction Kit (Qiagen). After assembly, this construct was transformed into *E. coli* One Shot™ TOP10 (Invitrogen). The recombinant gene insertion was confirmed by sequencing.

Sf9 cells (Novagen) were used to generate baculovirus stocks using the Bac-to-Bac system (Life Technologies) with a combination of bacmid donor cell DH10EmBacY (Geneva-Biotech), according to the manufacturer’s instructions. The titter of baculovirus stock was quantified using expressed YFP via fluorescence which serves as an infection marker.

Protein expression was carried out in sterile Vitlab Reagent bottles (VITLAB #100389) (37). Screw caps with a drilled hole were covered in gas-permeable non-woven adhesive tape (Batist^®^ Medical, #1320103112). Baculoviral P3 stock was used to infect Sf9 insect cells at 1.5 × 10^6^ cells·mL^-1^ in Insect-XPRESS media (Lonza #12-730Q) supplemented with 2% FBS (Capricorn #FBS-12A). For typical 1L expression, 3 mL of baculoviral stock was added to 1L of Sf9 cells. Infected cells were incubated in dark for 72 hours at 27 °C and 230 rpm (New Brunswick™ Innova® 42R), then the cells were harvested by centrifugation 1 500 g, 20 min, 4 °C. Pellet was washed in 40 mL of 1x PBS and centrifugated in a swinging-bucket rotor 3 000 g, 10 min, 4 °C. After centrifugation cells were snap-frozen in liquid nitrogen and stored at −80 °C until protein purification. From the typical 1L expression, we have obtained 8-12 g of cells containing REST-FL.

All the following purification steps were carried out with precooled buffers and instruments. Additionally, all the following steps were done at 4 °C. The frozen pellet was thawed on ice and resuspended in the lysis buffer (50 mM Sodium Phosphate, 500 mM NaCl, 5 % glycerol, 0.1 % Triton X-100, pH 8.0) with the addition of a final concentration of 1 mM PMSF and 5 mM benzamidine, leupeptin 1.9 μg·mL^-1^ and pepstatin A 3.1 μg·mL^-1^.

Typically, 30 mL of lysis buffer was used for insect cells harvested from 1L of media. The cell suspension was transferred into stainless steel container cooled on ice and sonicated for 5 minutes with an amplitude 30%, 1-second pulse-on, 1 second pulse-off with a 60-second process time cycle with a 15-second pause for extra cooling (Misonix Qsonica S-4000). The lysate was cleared by centrifugation of 80 000 g, 1 hour, 4 °C, and filtered by 0.45 μm Sterivex™ Filter Units (Merck Millipore).

REST-FL was purified in two-step chromatography and carried out in a cold room at 4°C with a chromatography system (NGC™ Chromatography System Quest™ 10 Plus, Bio-Rad).

Ultra-high affinity chromatography column Im7 (TriAltus) was used to capture CL7-tagged protein. We used a 1 mL Im7 column and a flow rate of 0.35 mL·min^-1^. After loading, the column was washed with Im7 Wash buffer (50 mM Sodium Phosphate, 500 mM NaCl, 5 % glycerol, pH 8.0) until the absorbance at 280 nm dropped down and stabilized, followed by a second wash with storage buffer (50 mM Sodium Phosphate, 150 mM NaCl, 2.5 % glycerol, pH 8.0).

Cleavage was carried out by premixed SortaseA (SrtA) protease (recombinantly prepared in our laboratory, variant Ca^2+^ independent SortaseA mutant with enhanced activity: SortaseA-6xHis (38)) to a final concentration of 90 μg·mL^-1^ in storage buffer with the addition of DTT to the final concentration of 1 mM. SortaseA has a recognition site: LPETG and leaves no extra amino acid on the N-terminal part of the protein of interest.

The protease was loaded to the Im7 column in two steps divided by 20 min intermediate equilibration and followed by a two-hour incubation. The tag-free protein was eluted at the flow rate of 0.3 mL·min^-1^. After the first affinity purification step, we typically obtained 0.6 mg of protein mixture from 1L of insect cell expression, as shown in Figure 3a, Elu fraction.

Affinity chromatography with His-trap column (HisTrap™HP, Cytiva), 1 mL column volume, flow rate 0.5 mL·min^-1^, was used to separate protease from the sample. The protein fractions were eluted in 0-300 mM imidazole gradient in a buffer of 50 mM Sodium Phosphate, 150 mM NaCl, imidazole, 2.5 % glycerol, pH 8.0. REST-FL was eluted at 15 mM imidazole concentration – typically, we obtained 0.2 mg of REST-FL in 5 – 6 mL. The remaining SrtA protease was eluted at a 150 mM imidazole as shown in SDS-PAGE Figure 3a.

Collected samples from the first main peak were concentrated by ultrafiltration on 30K Amicon^®^ Ultra-0.5 Centrifugal Filters Units (Merck Millipore) in a fixed rotor 14 000 g, 4 °C, until the concentration of protein was higher than 1 mg·mL^-1^ and imidazole in the solution was less than 5 mM (sample was diluted by storage buffer). After the final concentration, REST-FL was frozen in liquid nitrogen and stored at −80 °C. The typical yield from 2L of insect cell culture was 0.05 mg of pure REST-FL (final purity is in Figure 4). Purification was carried out within one day to preserve protein activity.

### NRSE duplex preparation

NRSE sequence, 21bp DNA duplex was prepared by annealing a fluorescently labeled (Alexa Fluor 488) oligonucleotide 5’-TTCAGCACCATGGACAGCGCC-3’ with its complementary unlabeled strand (13,14). Both oligonucleotides were purchased from Eurofins Genomics. The substrate was purified on a Mono Q 5/50 GL column (GE Healthcare) in buffer 25 mM Tris-HCl, 50 mM NaCl, pH 8.0 with elution of salt gradient up to 1 M.

### Binding analysis using fluorescence anisotropy

The interactions of REST-FL or REST-N62 with NRSE sequence labeled by Alexa Fluor 488 were measured on a FluoroMax-4 spectrofluorometer (Horiba Jobin Yvon) at 25°C. Fluorescence anisotropy was measured at an excitation wavelength of 496 nm and an emission wavelength of 521 nm. The slit width (both excitation and emission) for all measurements was 8 nm and the integration time was 1 second. Incubation time before the measurement was 25 minutes.

The cuvette contained 1.4 ml of labeled 7.5 nM NRSE DNA in a FA buffer (50 mM sodium phosphate, 150 mM NaCl, pH 8.0). The protein was titrated into the solution in the cuvette and measured after a 1 min incubation at 25 °C. Fluorescence anisotropy at each titration step was measured three times. The dissociation constant K_D_ was determined from non-linear fitting of the binding data using equation: FA = FAmin + FAmax*((x+ DNAt+ KD -((x+DNAt+KD)^2- (4*x*DNAt))^0.5)/(2*DNAt)) derived for one-site binding of REST on DNA (39). Experimental data were visualized, and the fit was calculated using ORIGIN® 9.9.0.225 (OriginLab).

### Coimmunoprecipitation and western blotting

Human cells HEK293T (kindly provided by Agnel Sfeir, MSK, USA) were cultured in DMEM (Gibco #41966) supplemented with 10% FBS (Biosera #FB-1090), 4 mM L-glutamine (Gibco #25030), 1x non-essential amino acids (Gibco #11140), 100 units·mL^-1^ penicillin and 100 μg·mL^-1^ streptomycin (Sigma-Aldrich #P4333) at 37°C with 5% CO2 for no longer than 13 passages.

3xFLAG-tagged REST-N62 (pcDNA5 vector), HALO-tagged REST-FL (pHAGE2 vector), and empty 3xFLAG-tagged pcDNA5 vector were used to transiently co-express proteins in different combinations together, as shown in Figure 6. Transfection was carried out by polyethyleneimine – PEI in a six-well plate for 4 hours. The cells continued to grow for the next 48 hours and later were harvested, washed with 1x PBS. The cells were lysed in CoIP lysis buffer (50 mM Tris-HCl, 150 mM NaCl, 1 mM EDTA, 0.5% Triton X-100, pH 7.4, and freshly added 10 mM NaF) with protease inhibitor cocktail cOmplete tablets (Roche) for 45 minutes in a cold room (4 °C) on a rotator. The supernatant was separated by centrifugation 15 000 g, 20 minutes, 4 °C, and mixed with Anti-FLAG® M2 magnetic beads (Millipore) – typically 900 μL of supernatant and 30 μL of preequilibrated M2 beads.

Incubation was carried out in a cold room on a rotator for 2 hours. After incubation, beads were washed with 5×500 μL CoIP lysis buffer. To elute IP fractions, beads were incubated 1 hour on ice in 30 μL of CoIP elution buffer (50 mM Tris-HCl, 150 mM NaCl, 1 mM EDTA, pH 7.4) containing 0.2 mg·mL^-1^ 3xFLAG peptide (Sigma-Aldrich).

Samples were loaded on 10% SDS-PAGE gel, separated, and wet transferred to a membrane (Cytiva, Amersham™ Protran®). Western blots were carried out with two different antibody setups: (i) FLAG tag was incubated with monoclonal ANTI-FLAG® M2-Peroxidase (HRP) produced in mouse (Sigma-Aldrich #A8592) diluted 1:3 000, and (ii) HALO tag was incubated with primary Anti-HaloTag® Monoclonal Antibody (Promega #G9211) diluted 1:1 000 with secondary antibody Anti-Mouse IgG (Fc specific) HRP (Sigma-Aldrich #A0168) diluted 1:5 000. Primary antibodies were incubated in a cold room overnight. The secondary antibody was incubated for one hour at RT on a rotator. Detection was carried out with SuperSignal™ West Femto Maximum Sensitivity Substrate (Thermo Scientific™ #34095) and visualized on Fusion FX (Vilber).

### SDS-PAGE analyses

We employed sodium dodecyl sulfate–denaturing polyacrylamide gel electrophoresis to analyze the purity of proteins in each stage of purification. We used self-casted fixed-concentration polyacrylamide SDS gels containing 10% or 12% acrylamide:bis-acrylamide (37:1) for the separations of proteins with similar molecular weight. Proteins were separated on fixed-concentration SDS-PAGE at E = 10 V·cm^-1^.

We used gradient 4–20% Mini-PROTEAN® TGX™ Precast Protein Gels (BioRad) for separation of proteins with varied molecular range. SDS-PAGE with gradient gels was performed according to the manufacturer’s protocol (200 V Standard): 15 minutes at 30 mA, followed by 20 minutes at 25 mA. We used proteins included in the molecular weight marker PageRuler™ Prestained Protein Ladder (Thermo Fisher Scientific) as standards. If needed, proteins in gels were visualized by Bio-Safe™ Coomassie Stain (BioRad).

### Negative stain electron microscopy

Four microliters of 0.02 mg·mL^-1^ REST protein sample were loaded on a plasma-treated TEM grid supplemented with a homemade 12 nm thick continuous carbon layer. The sample was incubated for 30 seconds, washed once in a drop of 2% uranyl acetate (UA), and then stained in the second UA drop for 60 seconds. Images were collected using the 200 kV Talos F200C transmission electron microscope equipped with the 16M Ceta detector using SerialEM (40). The overall dose was 15 e·Å^-2^ per single image, and data were collected at the calibrated pixel size of 2.65 Å. Data analysis was carried out in CryoSparc (41).

The analyzed dataset comprised 324 micrographs. Particles from 20 micrographs were picked manually and used to generate references for template picking. Following the reference-free 2D classification, 9218 particles were selected to determine the REST structure. The CryoSparc *ab initio* routine was used to determine the initial model and further classify the data in 3D. Finally, 3540 particles were refined to produce the REST model at 18.5 Å global resolution.

## Conclusions

Our study provides new insights into the structure-function relationships of human REST as a master regulator of neural differentiation. We presented the electron microscopy structure of full-length REST, suggesting that its toroidal shape can accommodate DNA in the central hole.

We developed protocols for the preparation and subsequent characterization of human REST and its splicing isoform REST-N62 allowing further structural and functional studies *in vitro*. We quantified REST binding to the canonical DNA. We excluded the high-affinity interaction of full-length REST and splicing variant REST-N62. We discussed the implications of the new structural findings to known functions of REST.

## Acknowledgments and funding

The Czech Science Foundation (19-18226S to C.H.) has primarily supported this research. The research has been carried out with institutional support of the Ministry of Education, Youth and Sports of the Czech Republic (LTAUSA19024 to C.H.); Institute of Biophysics of the Czech Academy of Sciences (68081707 to P.V, M.S. and C.H.). We acknowledge CEITEC MU Cryo-electron microscopy and tomography core facility, Proteomics, and Biomolecular Interactions and Crystallization core facility of CIISB, Instruct-CZ Centre supported by MEYS CR (LM2018127). T.B. is supported by Brno Ph.D. Talent Scholarship - funded by Brno City Municipality. We thank Victoria Marini for her critical reading and editing of the manuscript.

## Data availability

3D model of REST at 18.5 Å resolution is stored in Electron Microscopy Data Bank (EMDB) database as EMD-15858.

## Abbreviations

aa: amino acid(s)
bp: base pair(s)
CD: circular dichroism
CoIP: coimmunoprecipitation
DLS: dynamic light scattering
EM: electron microscopy
IDRs: intrinsically disordered regions
kDa: kilodaltons
NRSE: neuron-restrictive silencer element
NRSF: neuron-restrictive silencer factor
PCNA: proliferating cell nuclear antigen
RD1/2: repressor domain 1/2
RE1: repressor element-1
REST: RE1-silencing transcription factor
REST-FL: full-length REST
REST-N62: isoform of REST (also known as REST4 or REST4-S3)
ZF: zinc finger

## References

1. Lambert, S.A., Jolma, A., Campitelli, L.F., Das, P.K., Yin, Y.M., Albu, M., Chen, X.T., Taipale, J., Hughes, T.R. and Weirauch, M.T. (2018) The Human Transcription Factors. Cell, 172, 650–665.

2. Schoenherr, C.J., Paquette, A.J. and Anderson, D.J. (1996) Identification of potential target genes for the neuron-restrictive silencer factor. Proceedings of the National Academy of Sciences, 93, 9881–9886.

3. Chong, J.H.A., Tapiaramirez, J., Kim, S., Toledoaral, J.J., Zheng, Y.C., Boutros, M.C., Altshuller, Y.M., Frohman, M.A., Kraner, S.D. and Mandel, G. (1995) Rest - a Mammalian Silencer Protein That Restricts Sodium-Channel Gene-Expression to Neurons. Cell, 80, 949–957.

4. Schoenherr, C.J. and Anderson, D.J. (1995) The neuron-restrictive silencer factor (NRSF): a coordinate repressor of multiple neuron-specific genes. Science, 267, 1360–1363.

5. Lu, T., Aron, L., Zullo, J., Pan, Y., Kim, H., Chen, Y., Yang, T.H., Kim, H.M., Drake, D., Liu, X.S. et al. (2014) REST and stress resistance in ageing and Alzheimer’s disease. Nature, 507, 448–454.

6. Hwang, J.-Y. and Zukin, R.S. (2018) REST, a master transcriptional regulator in neurodegenerative disease. Current Opinion in Neurobiology, 48, 193–200.

7. Rodenas-Ruano, A., Chávez, A.E., Cossio, M.J., Castillo, P.E. and Zukin, R.S. (2012) REST-dependent epigenetic remodeling promotes the developmental switch in synaptic NMDA receptors. Nature Neuroscience, 15, 1382–1390.

8. Garcia-Manteiga, J.M., D’Alessandro, R. and Meldolesi, J. (2019) News about the Role of the Transcription Factor REST in Neurons: From Physiology to Pathology. Int J Mol Sci, 21.

9. Calderone, A., Jover, T., Noh, K.-m., Tanaka, H., Yokota, H., Lin, Y., Grooms, S.Y., Regis, R., Bennett, M.V.L. and Zukin, R.S. (2003) Ischemic Insults Derepress the Gene Silencer REST in Neurons Destined to Die. The Journal of Neuroscience, 23, 2112–2121.

10. McClelland, S., Flynn, C., Dubé, C., Richichi, C., Zha, Q., Ghestem, A., Esclapez, M., Bernard, C. and Baram, T.Z. (2011) Neuron-restrictive silencer factor-mediated hyperpolarization-activated cyclic nucleotide gated channelopathy in experimental temporal lobe epilepsy. Annals of Neurology, 70, 454–465.

11. Zuccato, C., Tartari, M., Crotti, A., Goffredo, D., Valenza, M., Conti, L., Cataudella, T., Leavitt, B.R., Hayden, M.R., Timmusk, T. et al. (2003) Huntingtin interacts with REST/NRSF to modulate the transcription of NRSE-controlled neuronal genes. Nature genetics, 35, 76–83.

12. Bruce, A.W., Donaldson, I.J., Wood, I.C., Yerbury, S.A., Sadowski, M.I., Chapman, M., Göttgens, B. and Buckley, N.J. (2004) Genome-wide analysis of repressor element 1 silencing transcription factor/neuron-restrictive silencing factor (REST/NRSF) target genes. Proceedings of the National Academy of Sciences, 101, 10458–10463.

13. Tang, Y., Jia, Z., Xu, H., Da, L.-t. and Wu, Q. (2021) Mechanism of REST/NRSF regulation of clustered protocadherin α genes. Nucleic acids research, 49, 4506–4521.

14. Thiel, G., Ekici, M. and Rössler, O.G. (2015) RE-1 silencing transcription factor (REST): a regulator of neuronal development and neuronal/endocrine function. Cell and Tissue Research, 359, 99–109.

15. Roopra, A., Qazi, R., Schoenike, B., Daley, T.J. and Morrison, J.F. (2004) Localized Domains of G9a-Mediated Histone Methylation Are Required for Silencing of Neuronal Genes. Molecular cell, 14, 727–738.

16. Ooi, L., Belyaev, N.D., Miyake, K., Wood, I.C. and Buckley, N.J. (2006) BRG1 Chromatin Remodeling Activity Is Required for Efficient Chromatin Binding by Repressor Element 1-silencing Transcription Factor (REST) and Facilitates REST-mediated Repression*. Journal of Biological Chemistry, 281, 38974–38980.

17. Palm, K., Metsis, M. and Timmusk, T. (1999) Neuron-specific splicing of zinc finger transcription factor REST/NRSF/XBR is frequent in neuroblastomas and conserved in human, mouse and rat. Molecular Brain Research, 72, 30–39.

18. Shimojo, M., Paquette, A.J., Anderson, D.J. and Hersh, L.B. (1999) Protein kinase A regulates cholinergic gene expression in PC12 cells: REST4 silences the silencing activity of neuron-restrictive silencer factor/REST. Mol Cell Biol, 19, 6788–6795.

19. Raj, B., O’Hanlon, D., Vessey, John P., Pan, Q., Ray, D., Buckley, Noel J., Miller, Freda D. and Blencowe, Benjamin J. (2011) Cross-Regulation between an Alternative Splicing Activator and a Transcription Repressor Controls Neurogenesis. Molecular cell, 43, 843–850.

20. Li, C.L., Wang, Z.F., Tang, X.Y., Zeng, L., Fan, X.T. and Li, Z. (2017) Molecular mechanisms and potential prognostic effects of REST and REST4 in glioma (Review). Mol Med Rep, 16, 3707–3712.

21. Varadi, M., Anyango, S., Deshpande, M., Nair, S., Natassia, C., Yordanova, G., Yuan, D., Stroe, O., Wood, G., Laydon, A. et al. (2021) AlphaFold Protein Structure Database: massively expanding the structural coverage of protein-sequence space with high-accuracy models. Nucleic acids research, 50, D439–D444.

22. Jumper, J., Evans, R., Pritzel, A., Green, T., Figurnov, M., Ronneberger, O., Tunyasuvunakool, K., Bates, R., Žídek, A., Potapenko, A. et al. (2021) Highly accurate protein structure prediction with AlphaFold. Nature, 596, 583–589.

23. Westbrook, T.F., Hu, G., Ang, X.L., Mulligan, P., Pavlova, N.N., Liang, A., Leng, Y., Maehr, R., Shi, Y., Harper, J.W. et al. (2008) SCFβ-TRCP controls oncogenic transformation and neural differentiation through REST degradation. Nature, 452, 370.

24. Janovic, T., Stojaspal, M., Veverka, P., Horakova, D. and Hofr, C. (2019) Human Telomere Repeat Binding Factor TRF1 Replaces TRF2 Bound to Shelterin Core Hub TIN2 when TPP1 Is Absent. Journal of molecular biology, 431, 3289–3301.

25. Veverka, P., Janovic, T. and Hofr, C. (2019) Quantitative Biology of Human Shelterin and Telomerase: Searching for the Weakest Point. International Journal of Molecular Sciences, 20, 13.

26. Fernandez Garcia, M., Moore, C.D., Schulz, K.N., Alberto, O., Donague, G., Harrison, M.M., Zhu, H. and Zaret, K.S. (2019) Structural Features of Transcription Factors Associating with Nucleosome Binding. Molecular cell, 75, 921–932.e926.

27. Li, H., Doruker, P., Hu, G. and Bahar, I. (2020) Modulation of Toroidal Proteins Dynamics in Favor of Functional Mechanisms upon Ligand Binding. Biophysical journal, 118, 1782–1794.

28. Lee, N., Park, S.J., Haddad, G., Kim, D.-K., Park, S.-M., Park, S.K. and Choi, K.Y. (2016) Interactomic analysis of REST/NRSF and implications of its functional links with the transcription suppressor TRIM28 during neuronal differentiation. Scientific Reports, 6, 39049.

29. Grimes, J.A., Nielsen, S.J., Battaglioli, E., Miska, E.A., Speh, J.C., Berry, D.L., Atouf, F., Holdener, B.C., Mandel, G. and Kouzarides, T. (2000) The Co-repressor mSin3A Is a Functional Component of the REST-CoREST Repressor Complex*. Journal of Biological Chemistry, 275, 9461–9467.

30. Barrios, Á.P., Gómez, A.V., Sáez, J.E., Ciossani, G., Toffolo, E., Battaglioli, E., Mattevi, A. and Andrés, M.E. (2014) Differential Properties of Transcriptional Complexes Formed by the CoREST Family. Molecular and Cellular Biology, 34, 2760–2770.

31. Yang, M., Gocke, C.B., Luo, X., Borek, D., Tomchick, D.R., Machius, M., Otwinowski, Z. and Yu, H. (2006) Structural Basis for CoREST-Dependent Demethylation of Nucleosomes by the Human LSD1 Histone Demethylase. Molecular cell, 23, 377–387.

32. Shimizu, K. and Toh, H. (2009) Interaction between Intrinsically Disordered Proteins Frequently Occurs in a Human Protein–Protein Interaction Network. Journal of molecular biology, 392, 1253–1265.

33. Nomura, M., Uda-Tochio, H., Murai, K., Mori, N. and Nishimura, Y. (2005) The Neural Repressor NRSF/REST Binds the PAH1 Domain of the Sin3 Corepressor by Using its Distinct Short Hydrophobic Helix. Journal of molecular biology, 354, 903–915.

34. Moumné, L., Betuing, S. and Caboche, J. (2013) Multiple Aspects of Gene Dysregulation in Huntington’s Disease. Front Neurol, 4, 127.

35. Kaneko, N., Hwang, J.-Y., Gertner, M., Pontarelli, F. and Zukin, R.S. (2014) Casein Kinase 1 Suppresses Activation of REST in Insulted Hippocampal Neurons and Halts Ischemia-Induced Neuronal Death. The Journal of Neuroscience, 34, 6030–6039.

36. Zhang, P., Lathia, J.D., Flavahan, W.A., Rich, J.N. and Mattson, M.P. (2009) Squelching glioblastoma stem cells by targeting REST for proteasomal degradation. Trends in Neurosciences, 32, 559–565.

37. Rieffel, S., Roest, S., Klopp, J., Carnal, S., Marti, S., Gerhartz, B. and Shrestha, B. (2014) Insect cell culture in reagent bottles. MethodsX, 1, 155–161.

38. Jeong, H.J., Abhiraman, G.C., Story, C.M., Ingram, J.R. and Dougan, S.K. (2017) Generation of Ca2+-independent sortase A mutants with enhanced activity for protein and cell surface labeling. PloS one, 12, e0189068.

39. Goodrich, J.A. and Kugel, J.F. (2006) Binding and Kinetics for Molecular Biologists. Cold Spring Harbor Laboratory Press.

40. Mastronarde, D.N. (2005) Automated electron microscope tomography using robust prediction of specimen movements. Journal of Structural Biology, 152, 36–51.

41. Punjani, A., Rubinstein, J.L., Fleet, D.J. and Brubaker, M.A. (2017) cryoSPARC: algorithms for rapid unsupervised cryo-EM structure determination. Nat Methods, 14, 290–296.

