## Supplementary materials for "Electron microscopy reveals toroidal shape of master neuronal cell differentiator REST – RE1-Silencing Transcription factor"

### Supplementary methods

#### Dynamic light scattering (DLS)

The homogeneity of the protein samples was determined by dynamic light scattering (DLS) in the Delsa Max Core (Beckman Coulter). Prior to measurement, 5 mg·mL$^{-1}$ REST-N62 in buffer (50 mM sodium phosphate, 150 mM NaCl, 5 % glycerol, pH 8.0) were centrifuged at 12 000 g, RT for 8 min. Twenty scans of 10 s data acquisition were averaged to examine the sample homogeneity. Data were collected and processed by software provided by Beckman Coulter. The presence of aggregates was determined based on a regularization fit of obtained autocorrelation functions of scattered light. The data were evaluated as the qualitative analysis of the aggregate's presence (the presence of peaks of Rh > 10 nm) for intensity-based data.

#### Circular dichroism (CD)

Far-UV CD measurements were performed on a J-815 spectrometer (Jasco) at 20 °C in a 1-mm Quartz cuvette (Hellma Analytics). CD spectra of 0.2 mg·mL$^{-1}$ REST-N62 in buffer (20 mM Sodium Phosphate, 150 mM NaF, pH 8.0) were acquired in the wavelength range 195–250 nm with 1 nm step at a scanning speed of 100 nm·min$^{-1}$. Each spectrum represents an average of ten accumulations. Subsequently, the buffer signal was subtracted, and data were converted from circular dichroism units to mean residue molar ellipticity (MRE). The presence of secondary structural elements was evaluated using BestSel software (1).

### Supplementary figures

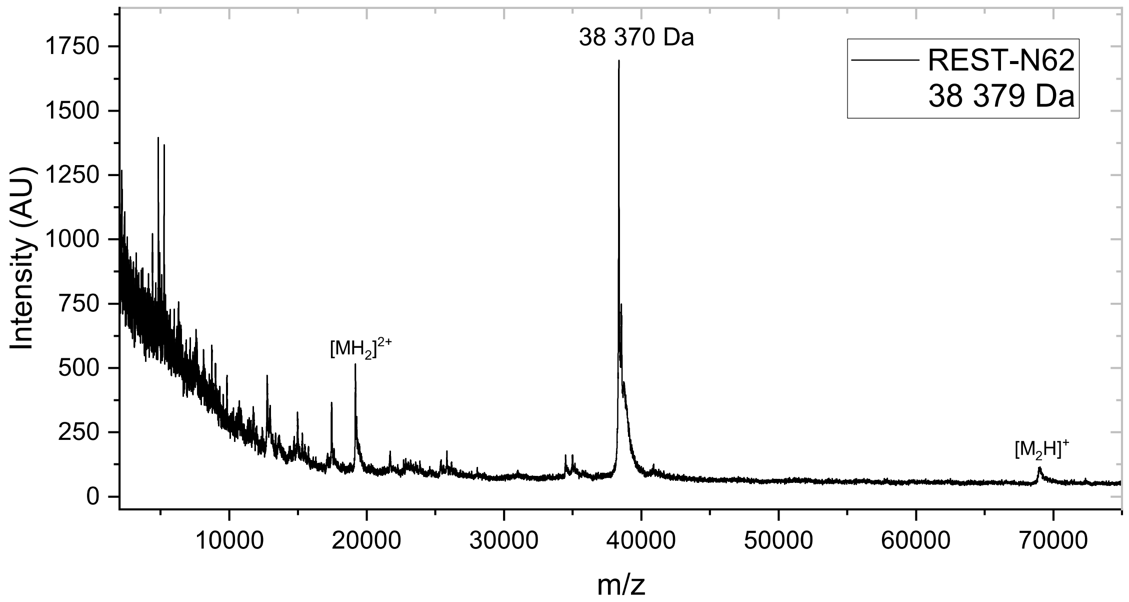


**Supplementary Figure S1: Mass Spectrometry confirms the identity of purified REST-N62.** MALDI-TOF Mass spectrometry measurement revealed the main peak with a molecular weight of 38 370 Da corresponding to the theoretical molecular weight of REST-N62 without tags (38 379 Da). All amino acids are present in purified REST-N62. The difference 9 Da between theoretical and measured molecular weight is probably caused by remaining salt ions during ionization.

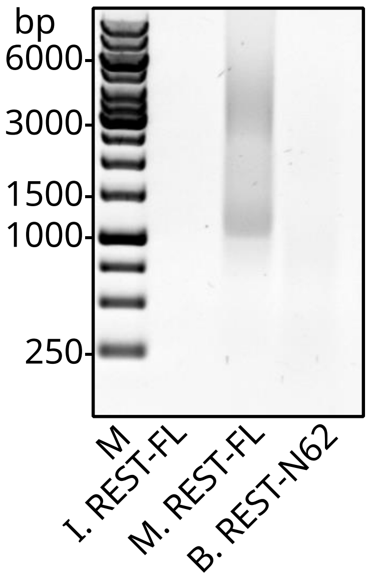


**Supplementary Figure S2: DNA content screening of purified protein constructs.** Protein samples were analyzed by electrophoresis using 1% agarose gel stained in ethidium bromide solution. Lane M - DNA marker GeneRuler 1 kb DNA Ladder. Lane I. REST-FL – protein purified from insect cells. Lane M. REST-FL – protein purified from mammalian cells. Lane B. REST-N62 – shortened splicing variant purified from *E. coli*.

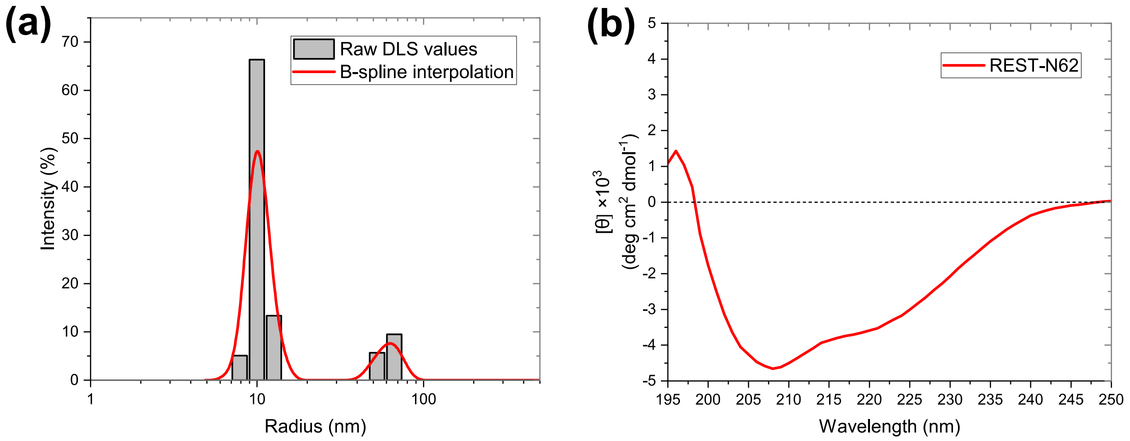


**Supplementary Figure S3: Homogeneity and secondary structure analyses of REST-N62.** (a) Dynamic light scattering (DLS) reveals a homogenous sample without significant aggregations. Minor content of a chaperone in the sample may contribute to the observed signal at around a 60 nm radius. The gray bar graph represents raw DLS values. The red line represents B-spline interpolated raw data. (b) The circular dichroism (CD) far-UV spectrum shows a profile corresponding to mainly the alpha-helical structure of the purified REST-N62.

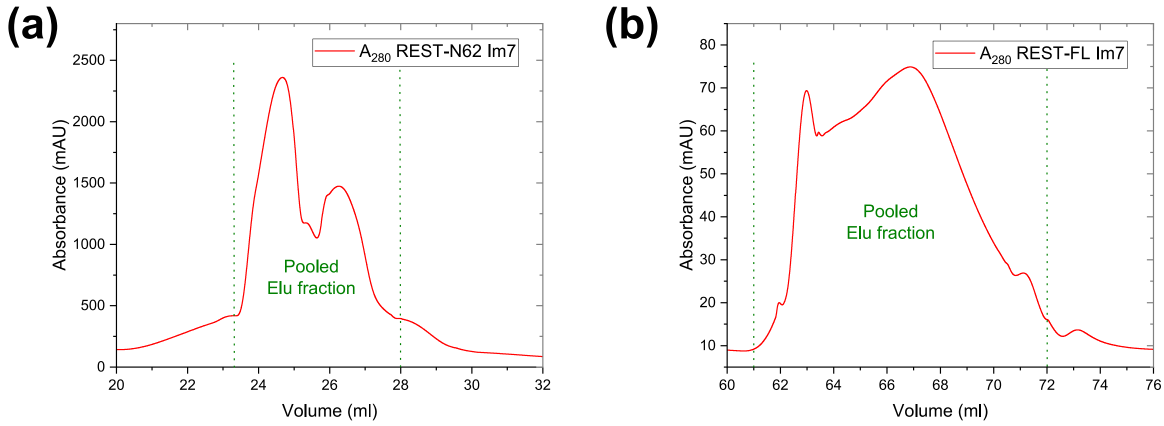


**Supplementary Figure S4: Ultra affinity Im7 column chromatograms.** (a) REST-N62 and (b) REST-FL chromatographic traces of affinity purification using the Im7-CL7 system. Elu fraction – pooled fractions after release from the Im7 column. Green marks denominate collection ranges. A double-peak appearance of chromatographic traces occurs as a result of 20-minute incubation with a protease that releases the untagged protein construct.

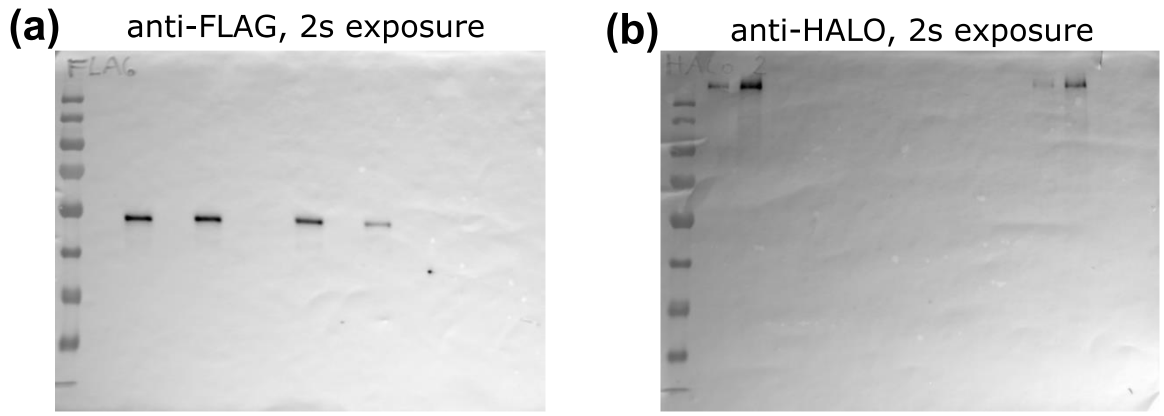


**Supplementary Figure S5: Original images of western blots used for Coimmunoprecipitation of REST-FL and REST-N62.** Both membranes had the same samples in gels' wells: Marker, 4x whole cell extract (input fraction), one empty well, 4x bound fraction, one empty well, and 4x unbound fraction. (a) Western blot of anti-FLAG with 2 seconds of exposure. (b) Western blot of anti-HALO with 2 seconds of exposure.

### Supplementary references

1. Micsonai, A., Moussong, É., Wien, F., Boros, E., Vadászi, H., Murvai, N., Lee, Y.-H., Molnár, T., Réfrégiers, M., Goto, Y. *et al.* (2022) BeStSel: webserver for secondary structure and fold prediction for protein CD spectroscopy. *Nucleic acids research*, **50**, W90-W98.
